# Microbial load on hair tools of tertiary-level students in Ghana; A case study of the University of Science and Technology, Ghana

**DOI:** 10.64898/2026.08.09.743799

**Authors:** Gyasi Junior Darko, Helena Addison, Angela Baaba Forson, Michael Nkrumah-Appau, William Gariba Akanwariwiak

**Author notes:** **Corresponding Author:** Gyasi Junior Darko, Kumasi Centre for Collaborative Research in Tropical Medicine, Ghana, Kwame Nkrumah University of Science & Technology, Kumasi - Ghana. **Email Addresses:** Helena Addison, Angela Baaba Forson, Michael Nkrumah-Appau, William Gariba Akanwariwiak. Author Contributions: WGA was the initiator of the study. WGA, GJD, HA, ABF together conceptualized and developed the study. GJD, HA, ABF and MNA conducted the laboratory research. MNA conducted data analyses for the study. All authors drafted, read, edited and gave approval for the final manuscript.

## Abstract

**Background:** Hair tools such as hairbrushes and combs allow for various styling options to produce desired hairstyles among various people. However, there is the risk of they serving as fomites for infection and contamination especially among people in close habitation.

**Objectives:** This study therefore decided to investigate the trend of microbial populations on these hair grooming tools in universities such as KNUST to inform student hygiene practices and disease prevention strategies.

**Methods:** 30 students were randomly selected for the study, and swab samples from different hairbrushes and combs were taken for microbial investigation. Microbial isolates were identified based on their morphological and biochemical characteristics. Determination of efficacy of different cleaning methods for hair tools was also done.

**Results:** The study found an average bacterial and fungal count of 7.4×10^2^ CFU/ml and 4.6×10^3^ CFU/ml, respectively. The bacterial isolates suspected included *Staphylococcus aureus, Staphylococcus epidermidis, Streptococcus sp*., *Bacillus subtilis*, and *Corynebacterium sp*. The fungal isolates included *Aspergillus* species, *Penicillium sp*., *Rhizopus sp*., *Neurospora sp*., *Colletotrichum gloeosporioides* and *Curlvularia sp*. Correlation analysis showed higher bacterial numbers significantly associated with the presence of hair diseases such as dandruff (p=0.046). Water and detergent were found to be the most effective method of eliminating microbial content from hairbrushes and combs.

**Conclusion:** This study uncovered a variety of microbes on KNUST students’ combs and hairbrushes, which is evident of microbial contamination. While these numbers are relatively low, this study highlights the need for students to still follow good hygiene procedures and implement efficient cleaning techniques of hair tools, as they may still serve as an ideal environment to harbor and transfer microbes

## INTRODUCTION

The usage of hair grooming tools predates modern civilizations as history have left behind traces of their use with hieroglyphs and artifacts from thousand years ago, for example, show individuals brushing their hair with brushes made of wood, bone, or ivory from elephant tusks (Termix, 2020). Hair grooming tools such as hairbrushes and combs have a lot of significance in personal grooming routines as they contribute to both practical, hygiene and aesthetic purposes (Meadow et al., 2015; Type, 2015).

These tools are used on daily basis to manage and groom hair, as well as to keep hygiene. However, frequency and profoundity of their use may gather skin cells, hair debris, product residues, and germs, which could result in the formation of microbial reservoirs; particularly bacteria and fungi (Sanke, 2022). There is also the concern of how these harbored microbes can aid in the spread of infection and contamination. These microbes can be transferred from the hair scalp to personal grooming tools like hairbrushes and hair combs.

Bacteria are among the most prevalent microbes detected on used hairbrushes and combs (Edward et al., 2015) including both Gram-positive and Gram-negative species (Iuka et al., 2014). *Pseudomonas aeruginosa*, which can cause infections of the skin and soft tissues (Nagoba et al., 2017), and *Staphylococcus aureus*, a bacterium linked to skin infections and drug resistance (Wilkinson et al., 2024), are common bacterial contamination on grooming products (deviS, 2021).Other bacteria that are frequently discovered on combs and hairbrushes include species of *Corynebacterium, Bacillus, and Streptococcus*.

Used hairbrushes and combs are also often covered in fungi, such as yeasts and molds (Alharbi & Alhashim, 2021). These microbes can colonize grooming instruments, especially if they are not properly cleaned and dried after each use. Species of Candida, Aspergillus, Trichophyton are all common fungi found on grooming items (Edward et al., 2015).

A few studies have also identified viral infection on worn combs and hairbrushes (Alharbi & Alhashim, 2021) although, they are less well studied than bacteria and fungus. Human papillomavirus (HPV) and (HSV) herpes simplex virus are the common of viral pollutants implicated (Avci & Ertam, 2014). Although direct touch or respiratory droplets are the most typical ways for viruses to spread, studies have found viral DNA or RNA on contaminated surfaces, such as combs and hairbrushes (Nandini Shetty, Julian W Tang, 2009).

Humans living in close proximity of one another are often at highest risk of person-to-person contamination/infection (Meadow et al., 2015). The factors that drive this risk include population volume, human contact, inter-use of personal effects (e.g., bath items, culirinary items, bodycare tools, etc.), and body fluids (Meadow et al., 2015).

University campuses are one such places where hygiene is an issue of concern due to the closeness of students to one another in their habitations; and some studies have shown that university students’ hygiene practices and the cleanliness of their grooming supplies may be impacted by shared living arrangements, public grooming locations, and hectic class schedules (Kabir et al., 2021).

Prior research in this field of microbiology has predominantly focused on broad hygiene habits or the microbiological contamination of various surfaces (Vandini et al., 2014), resulting in a knowledge gap regarding the particular hygiene practices and microbiological profiles linked to personal grooming items. The absence of extensive research on the types and aggregate of microbes on used hairbrushes and hair combs, particularly among student populations, worsens the problem. An inadequate understanding of the microbiological effects of personal hygiene practices as well as less knowledge on the effectiveness of various cleaning methods can result in poor hygiene habits and higher health risks. Closing this information gap will thus be instrumental in mitigating possible health risks related to grooming equipment functioning as fomites for microbial contamination, promote educated hygiene habits, and protect the health and wellbeing of students, especially in tertiary level education such as KNUST.

Specifically, this study sought to: identify the microorganisms isolated from used hairbrushes and combs among students and determine the microbial load on used hairbrushes and combs among students.

## MATERIALS AND METHODS

### Study design and setting

This was a cross-sectional study that establishes the diversity of the microbial composition found on used hairbrushes and combs of students on Kwame Nkrumah University of Science and Technology (KNUST) Campus. The university has the second largest student population in the country, currently of about 85,000 according to University Information Technology Services (UITS), KNUST and covers a land area of 2,512.96 acres.

### Study population and size

A total of 30 undergraduate students of the university were included in this study.

### Eligibility criteria

Only undergraduate students who consented to participate in the study were included

### Study procedure

#### Demographic & history data collection

A participant information leaflet (PIL) and attached structured questionnaire developed in English (Appendix I) was distributed to participants. The PIL sought participant consent and provided assurance of anonymity. Each questionnaire was assigned a distinct participant identification number (PIDN) and collated data including participant sex, age, academic level, knowledge of personal hygiene, hair tools maintenance and history of hair infection.

### Sample collection

Swabs were taken from hairbrushes and combs using sterilized swab sticks moistened in peptone water. Collected samples were stored temporarily on ice at 4-7°C at the sample site and then transported to the microbiology laboratory for bacteriological and mycological investigations.

### Laboratory investigations

#### Microbial culture and isolation

Serial dilutions of sample solutions were prepared up to 1 x 10^-4^ concentrations and were plated on Mannitol Salt Agar (MSA) and Potato Dextrose Agar and incubated for 18-24 hours and 2-5 days at 37°C; for bacteria and fungi respectively.

After culture, bacteria isolated from MSA plates were identified and enumerated using a combination of colonial morphology, a colony counter and Gram’s staining.

#### Confirmatory tests for Staphylococcus sp

Citrate and catalase tests were conducted to detect and select for Staphylococcus sp. present. The catalase test was to distinguish Staphylococcus sp. from Streptococcus sp. based on their specific ability to produce the enzyme catalase while the citrate test was to identify the target organism based on their possession of the enzyme citrate which is distinct to *Staphylococcus sp* (Dcd, 2007; Reiner, 2013).

### Statistical inferences

Data was entered in Microsoft Excel Professional Plus 2024 Preview and was statistically analyzed using Stata 15. Paired t-tests were used to compare means of continuous variables while the chi-square test of independence (t-test with measures of association) was conducted to determine the possible association between hygiene practices of participants and microbial contamination. A p-value less than 0.05 was considered statistically significant. Graphs and charts were created using GraphPad Prism version 9.5.1 (GraphPad Software, San Diego, CA, USA).

## RESULTS

### Demography of participants

Thirty (30) individuals were randomly selected and interviewed for this study between May and July 2024. Participants were between the ages of 17-25 years (mean = 20.3) and were equally distributed between males and females. Majority of the participants (40.0%) were in their first year and from the College of Humanities and Social Sciences (Table 1).

**Table 1:** Demography of participants.

| Characteristic | Value | Frequency (n) | Percentage (%) |
| --- | --- | --- | --- |
| Gender | Male | 15 | 50.0 |
|  | Female | 15 | 50.0 |
| Academic level | 100 | 12 | 40.0 |
|  | 200 | 8 | 26.7 |
|  | 300 | 1 | 3.3 |
|  | 400 | 9 | 30.0 |
| Age<br>(mean = 20.3) | 17 | 1 | 3.3 |
|  | 18 | 3 | 10.0 |
|  | 19 | 9 | 30.0 |
|  | 20 | 5 | 16.7 |
|  | 21 | 5 | 16.7 |
|  | 22 | 3 | 10.0 |
|  | 23 | 2 | 6.7 |
|  | 24 | 1 | 3.3 |
|  | 25 | 1 | 3.3 |
| College of affiliation | Science | 7 | 23.3 |
|  | Agriculture & Natural resources | 1 | 3.3 |
|  | Arts & Built Environment | 2 | 6.7 |
|  | Engineering | 9 | 30.0 |
|  | Humanities & Social Sciences | 11 | 36.7 |

### Knowledge of microbiology of hair and hygiene

Of the 30 participants, 70.0% participants responded as having some knowledge of microbes (Figure 1) with 63.3 % knowing of microbes being present in hair (Figure 2).

**Figure 1:**
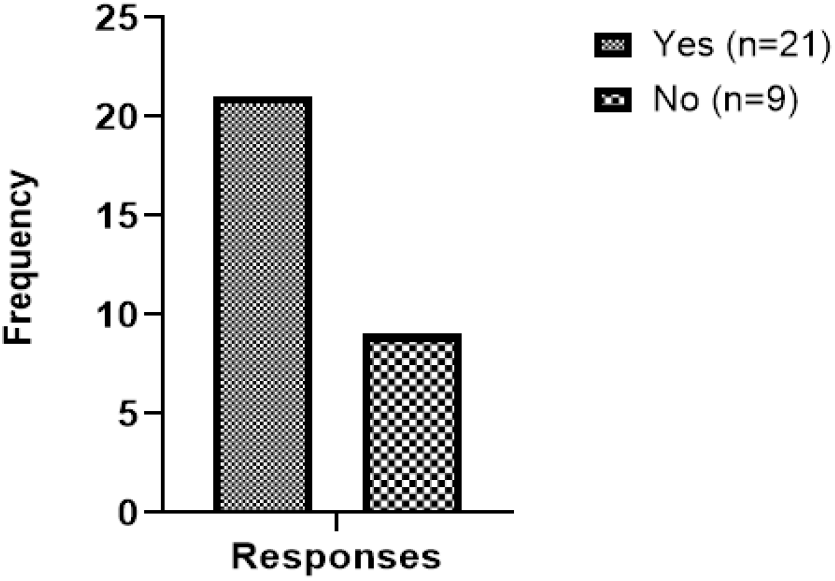
Knowledge of microbes

**Figure 2:**
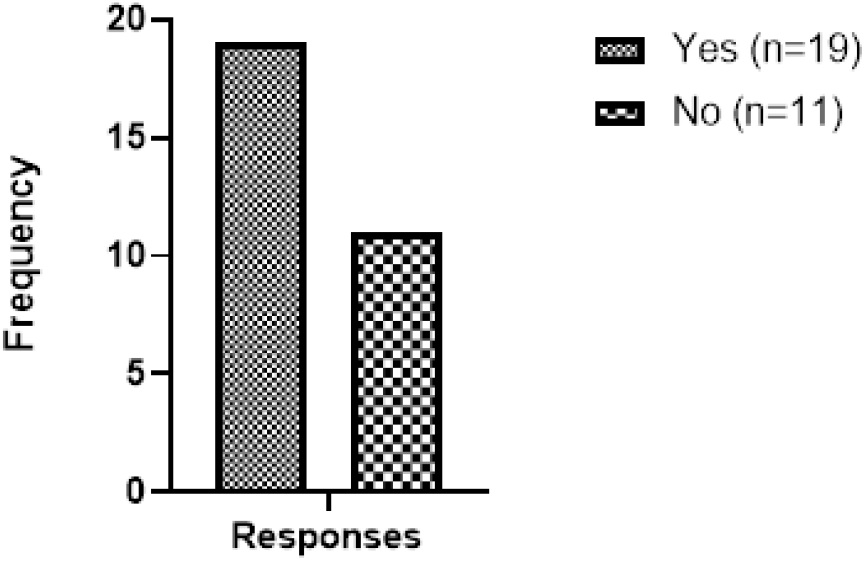
Knowledge of microbes in hair

### Hair and hair tools hygiene practices

The study found that majority of students (70.0%) shared hair tools with others (Figure 3) while less than half (46.6%) had used their hair tools beyond 6 months (Figure 4). Only 23.4% of participants reported having any history of hair diseases. With respect to hygiene of hair tools, more than half (63.3%) of participants cleaned their tools regularly (Figure 5) with only 26.7% cleaning them with the aid of a disinfecting agent (Figure 6).

**Figure 3:**
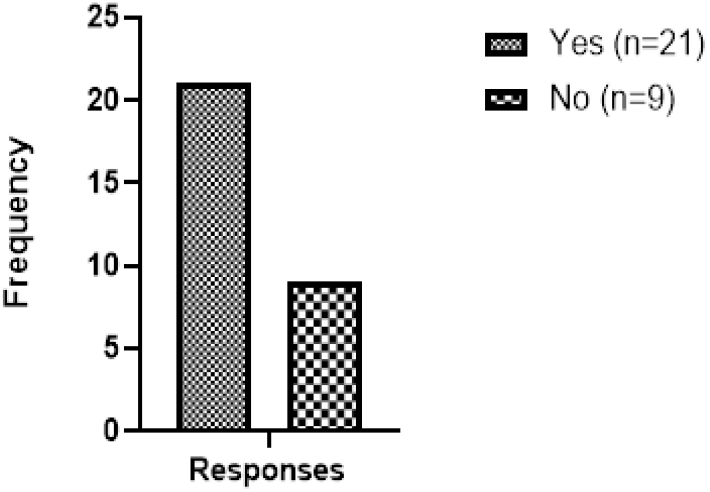
Response to sharing of hair tools

**Figure 4:**
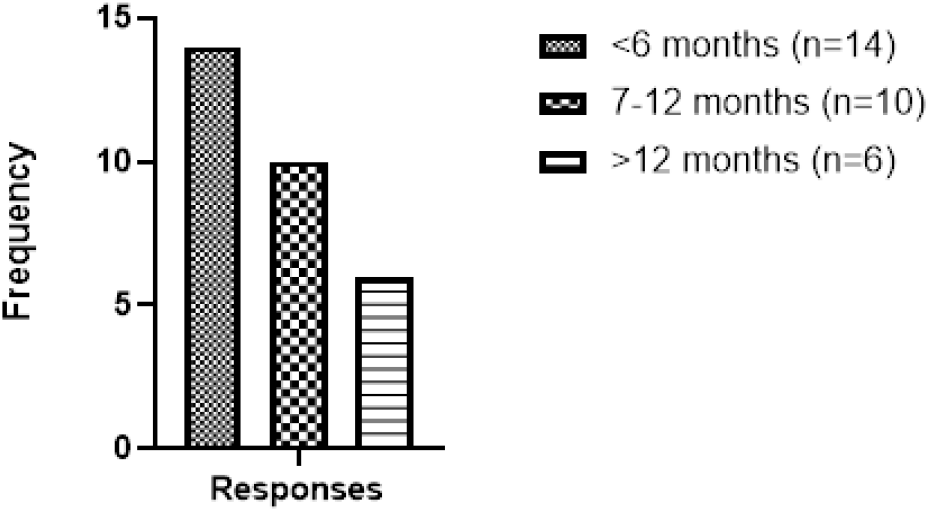
Duration of usage of hair tool/s

**Figure 5:**
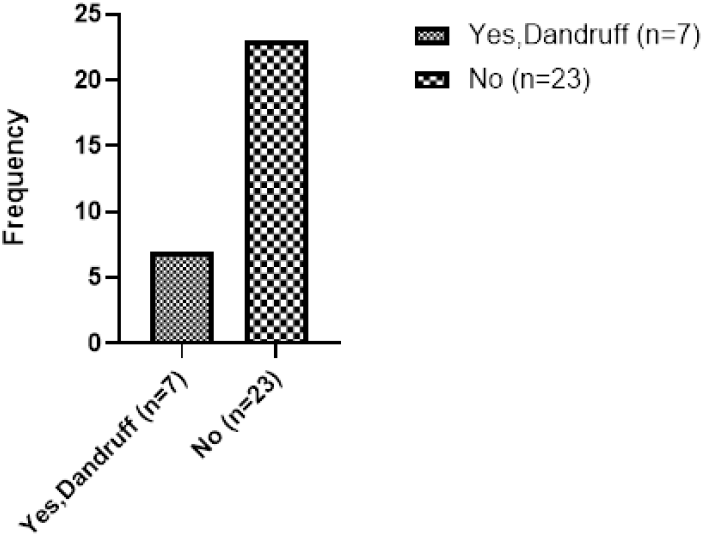
History of hair disease/s

**Figure 6:**
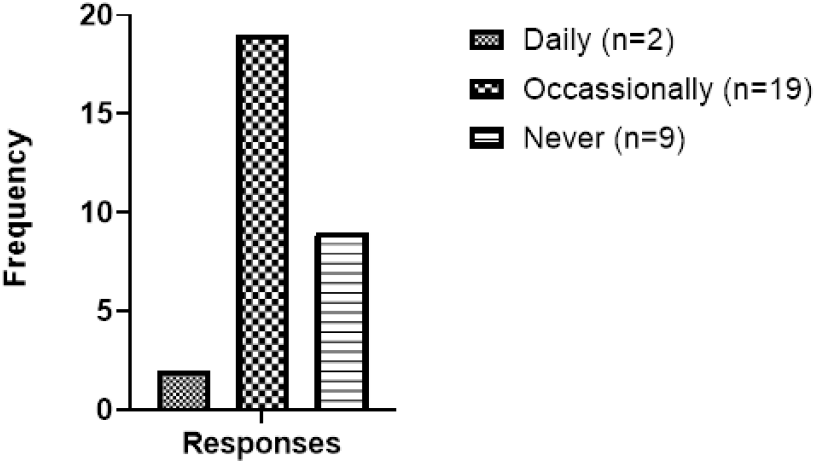
Frequency of cleaning of tools

**Figure 7:**
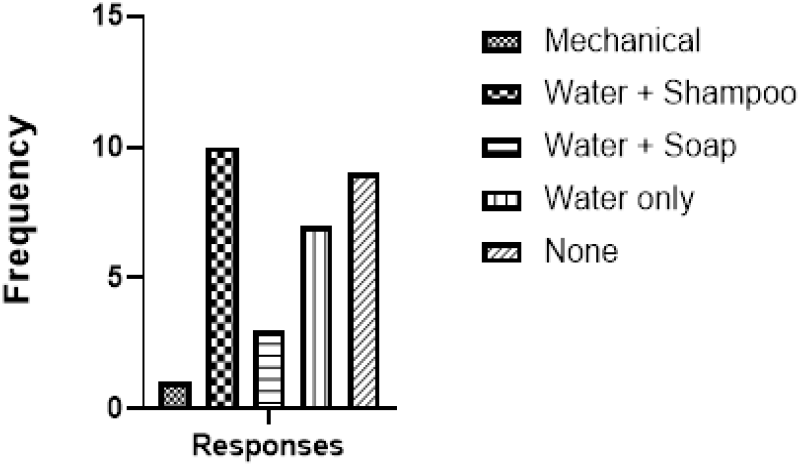
Method/s of cleaning

### Bacterial isolation from hair tools

A total of 35 bacterial colonies were isolated from the 30 collected samples with 19 (54.28 %) of the isolates being Gram-positive cocci indicating probable presence of *Staphylococcus sp*. (catalase-positive) and Streptococcus sp.(catalase-negative). The remaining 16 (45.7 %) isolates were Gram-positive rods. Catalase and citrate tests were conducted on the isolated GPC (19) isolates. This showed 17 (89.5%) positives for catalase and 15 (78.9%) positives for citrate test (Figure 8). These results are therefore showing a 48.6% prevalence of Staphylococcus sp. on hair tools of tertiary-level students.

**Figure 8:**
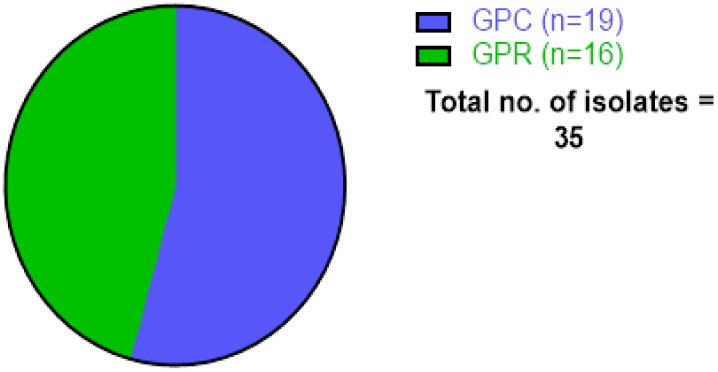
Gram reaction of isolated bacteria

**Figure 9:**
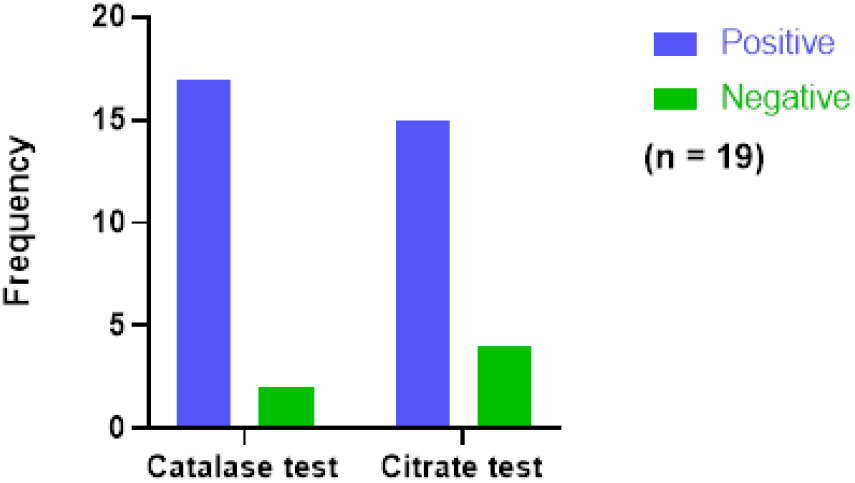
Biochemical confirmation of *Staphylococcus sp*.

### Fungal isolation from hair tools

Of samples from the 30 participants, 56 fungal isolates were identified from colonial morphology. *Aspergillus fumigatus* 22 (39.3 %) was the most commonly isolated among 8 other species identified; including *Aspergillus niger* (30.4 %), *Neurospora sp*. (10.7 %), *Colletotricum gloesporiodes* (9 %), *Aspergillus flavus* (3.6 %), *Penicillium sp*. (3.6 %), *Rhizopus sp*. (1.7 %) and *Curvularia sp*. (1.7 %) [Figure 10].

**Figure 10:**
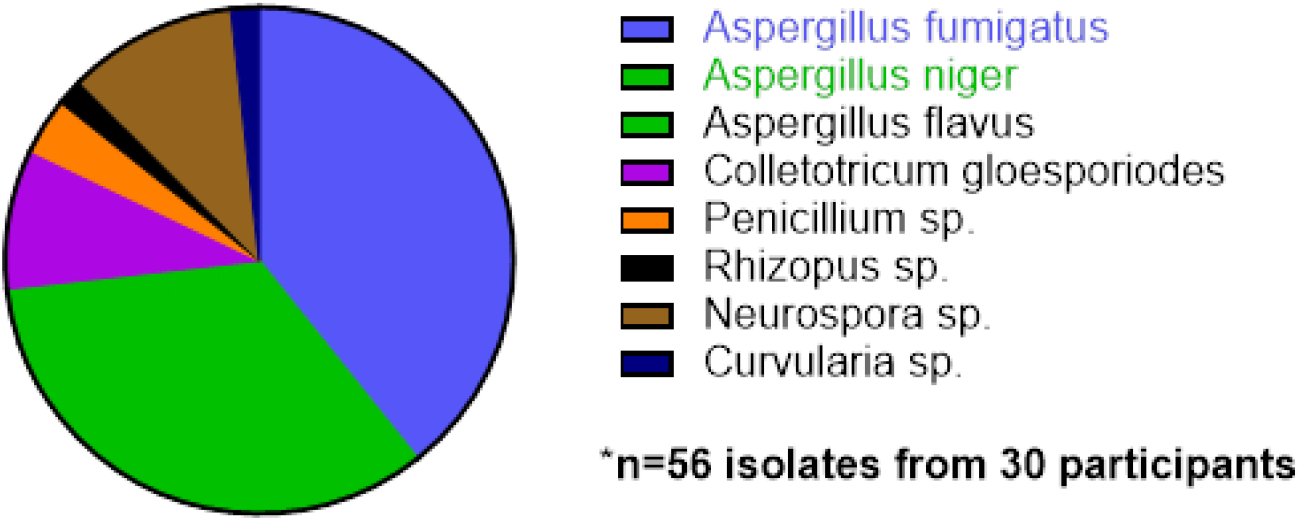
Isolated fungal species

### Microbial load on sampled hair tools

The average bacterial and fungal counts per positive sample were 4.5613×103cfu/*ml* and 7.353×102 cfu/ml respectively; with nominal ranges between 2.0 × 10^2^ cfu/ml - 7.0 × 10^3^ cfu/ml and fungal count ranges from (0 - 1.9×10^4^ cfu/ml respectively (Table 2).

**Table 2:** Microbial load on hair tools of student participants.

| Microbial group | Count (CFU/ml*) | Min. – Max. (CFU/ml*) |
| --- | --- | --- |
| Bacterial count | 7.353 x 10<sup>2</sup> | 2.0 x 10<sup>1</sup> – 7.0 x 10<sup>3</sup> |
| Fungal count | 4.5613 x 10<sup>3</sup> | 0 – 1.9 x 10<sup>4</sup> |

\*CFU/ml = coliform forming unit per millilitre

### Relationship between bacterial contamination of hair tools and hygiene practices

Using a ranking system of bacterial counts of <30 CFU/ml as low/negligible, 31 – 300 CFU/ml as high and >300 CFU/ml as very high, the study found relationships between hygiene practices such as frequent cleaning of hair tools and presence/history of hair disease to be associated with higher numbers of bacteria on hair tools. This relationship was especially significant for individuals who had histories of hair disease (Table 3).

**Table 3:** Association between hair hygiene practices and bacterial contamination of hair tools.

| Hair hygiene practices | Range of microbial contamination |  |  | p-value |
| --- | --- | --- | --- | --- |
|  | Low<br>(n=3) | High<br>(n=7) | Very high<br>(n=20) |  |
| Sharing hair tools (Yes/No) | 0/3 | 3/4 | 6/14 | 0.399 |
| Duration of use of current hair tool(s)<br>(<6mth/6-12mth/>12mth) | 1/1/1 | 4/1/2 | 9/8/3 | 0.719 |
| Frequency of use of hair product(s)<br>(Always/Never/Sometimes) | 0/0/3 | 2/2/4 | 2/5/13 | 0.448 |
| Presence of hair disease (Dandruff)<br>(Yes/No) | 3/0 | 3/4 | 17/3 | **0.046** |
| Cleaning of hair tools | 2/1 | 1/6 | 10/10 | 0.180 |
| Frequency of cleaning | 0/1/2 | 1/1/5 | 1/7/12 | 0.780 |

*P<0.05 was considered to be of statistical significance*

### Effect of different cleaning methods on reducing microbial load on hair tools

Results from the study highlighted the effectiveness of various hair tool cleaning methods on reduction of microbial population as a guide to advice on hygiene. Results showed that including chemical cleaning agents with water increased the efficiency in reducing microbial populations. Washing with water only removed 3.2% of bacteria which was poor in performance compared with including antiseptic (42.9%), soap (90.6%) and detergent (97.7%) [Table 4].

**Table 4:** Cleaning methods and bacterial numbers on hair tools.

| Cleaning method | Bacterial count (CFU/ml) |  |  | p-value<br>(signed-rank<br>test) |
| --- | --- | --- | --- | --- |
|  | Before cleaning | After cleaning | Difference/ % |  |
| Water only | $2.21 \times 10^2$ | $2.14 \times 10^2$ | $7.0 \times 10^1$ (3.2) | **\*0.0679** |
| Water + Antiseptic | $4.9 \times 10^1$ | $2.8 \times 10^1$ | $2.1 \times 10^1$ (42.9) | |
| Water + Soap | $9.7 \times 10^2$ | $9.1 \times 10^1$ | $6.0 \times 10^2$ (90.6) | |
| Water + detergent | $1.38 \times 10^2$ | $3.1 \times 10^1$ | $1.07 \times 10^2$ (97.7) | |

\**p-value* >0.05 ∴ differences not statistically significant

## DISCUSSION

Hair and hair tool hygiene is of particular importance in settings where individuals share close proximity due to their propensity to serve as viable fomites and reservoirs of infectious pathogens, their resistance genes, and as well as their spread (Wilkie et al., 2024). Previous studies also showed how this hair tools do indeed harbor significant numbers of bacteria and even fungi. For example, Ebuara et al., (2020) who analyzed samples from clippers, combs and hairbrushes from 40 different beauty salons in Taraba State, Nigeria. They also found pathogenic bacteria, ranging from *Staphylococcus sp*., *Bacillus sp*., and Streptococcus, as well as pathogenic fungi of the *Aspergillus, Trichophyton, Malasseza, Mucor*, and *Microsporum* genera. The findings of our study ascertain these observations and provide caution on the knowledge of these tools being fomites of possible microbial transmission.

Staphylococcus sp. particularly is a bacterial group of human and even environmental significance(Cuny et al., 2024), especially in the era of antimicrobial resistance and in the general fabric of infection prevention and control (Sigudu et al., 2024). They have been implicated in a number of opportunistic and active infections including psoriasis and cellulitis (Ng et al., 2017). The study therefore identifying significant numbers of *Staphylococcus sp*. therefore becomes of concern as although they are mainly commensal of the dermis (Battaglia & Garrett-Sinha, 2023), they may still serve as carriers and possible implicated pathogens in immunosuppressed individuals as well as even when they find themselves at unnatural anatomical sites (Sasson et al., 2017)

An important component of hair hygiene, as indicative of the study, is ensuring the hygiene of hair tools by cleaning them. Findings of the study also show that cleaning with water only may not be sufficient for effective bacterial removal and is better when supplemented with chemical-based bacteria-removal agents like soap, antiseptics and detergents. These agents often contain antimicrobial agents, like phenolic compounds, chlorine-based agents, or quaternary ammonium compounds (quats), which make them more efficient at lowering the number of microorganisms on surfaces (Fraise et al., 2004). Some, like most soaps, function by altering cell membranes of bacteria thus destroying them to remove them more effectively (Arellano et al., 2023). It is however notable that this depends on the type of soap used as not all may be impregnated with substances of antimicrobial properties (Ungphaiboon et al., 2005). Ideal soaps recommended should contain antimicrobial agents that inhibit the bacterial growth and even kill them (Kim & Rhee, 2016).

## CONCLUSION

The study realizes the significance of hair tools in the daily lives of people; however, if not properly monitored, these tools could become active fomites for microbial transmission among populations. Almost 1 in every 2 hair tools being used by tertiary-level students shows the presence of *Staphylococcus sp*.; which are organisms often implicated in infectious cases. Significant numbers of fungi species were also identified. These are all indicative of the need for particulate care for hair care tools which is often neglected; considering their potential for facilitating microbial spread. Especially in close-settlement settings like tertiary campuses, this is of prime essence. The study also concludes that hair tool hygiene practices such as frequent washing could be very significant in controlling microbial numbers, ergo, the possibilities of transmission. The washing should however be done with the use of chemical agents such as soap or detergents as much as possible for improved efficiency.

## REFERENCES

1. Alharbi, N., & Alhashim, H. M. (2021). Identification of Pathogenic Microbes in Tools of Beauty Salon in Jeddah City. Biosciences Biotechnology Research Asia, 18(4), 743–756. 10.13005/bbra/2956

2. Arellano, H., Nardello-Rataj, V., Szunerits, S., Boukherroub, R., & Fameau, A.-L. (2023). Saturated long chain fatty acids as possible natural alternative antibacterial agents: Opportunities and challenges. Advances in Colloid and Interface Science, 318, 102952. 10.1016/j.cis.2023.102952

3. Avci, O., & Ertam, I. (2014). Viral infections of the face. Clinics in Dermatology, 32(6), 715–733. 10.1016/j.clindermatol.2014.02.010

4. Battaglia, M., & Garrett-Sinha, L. A. (2023). Staphylococcus xylosus and Staphylococcus aureus as commensals and pathogens on murine skin. Laboratory Animal Research, 39(1), 1–13. 10.1186/s42826-023-00169-0

5. Cuny, C., Layer-Nicolaou, F., Werner, G., & Witte, W. (2024). A look at staphylococci from the one health perspective. International Journal of Medical Microbiology, 314, 151604. 10.1016/j.ijmm.2024.151604

6. Dcd, R. (2007). Chapter 12 Chapter 12. 1905(page 114), 46–48.

7. deviS, R. (2021). Evaluation of Microbial Load on Combs in Various Places Within the Tirupur District, Tamilnadu, and Sensitivity Pattern of Disinfectants. 9(8), 774–786. http://www.ijcrt.org

8. Ebuara, F. U., Imarenezor, E. P. K., Brown, S. T. C., Aso, R. E., Obasi, B. C., & Tyovenda, E. T. (2020). Isolation and Identification of Pathogenic Microorganisms Associated With Barbers’ Equipment in Wukari, Taraba State, Nigeria. FUW Trends in Science & Technology Journal, http://Www.Ftstjournal.Come-ISSN, 5(1), 215–218. http://www.ftstjournal.com

9. Edward, S. M., Megantara, I., & Dwiyana, R. F. (2015). Detection of Fungi in Hair-brushes in Beauty Salons at Jatinangor. Althea Medical Journal, 2(4), 516–520. 10.15850/amj.v2n4.636

10. Fraise, A. P., Lambert, P. A., & Maillard, J.-Y. (Eds.). (2004). Russell, Hugo & Ayliffe’s Principles and Practice of Disinfection, Preservation & Sterilization. Blackwell Publishing Ltd. 10.1002/9780470755884

11. Iuka, M., Stanley, C., Ifeanyi, O. E., Chinedum, O. K., & Onyekachi, I. S. (2014). Evaluation of Microbial Contamination of Tools Used In Hair Dressing Salons in Michael Okpara. IOSR Journal of Dental and Medical Sciences (IOSR-JDMS) e-ISSN, 13(7), 22–27. http://www.iosrjournals.org

12. Kabir, A., Roy, S., Begum, K., Kabir, A. H., & Miah, M. S. (2021). Factors influencing sanitation and hygiene practices among students in a public university in Bangladesh. PLOS ONE, 16(9), e0257663. 10.1371/journal.pone.0257663

13. Kim, S. A., & Rhee, M. S. (2016). Microbicidal effects of plain soap vs triclocarban-based antibacterial soap. Journal of Hospital Infection, 94(3), 276–280. 10.1016/j.jhin.2016.07.010

14. Meadow, J. F., Altrichter, A. E., Bateman, A. C., Stenson, J., Brown, G. Z., Green, J. L., & Bohannan, B. J. M. (2015). Humans differ in their personal microbial cloud. PeerJ, 2015(9), 1–22. 10.7717/peerj.1258

15. Nagoba, B., Davane, M., Gandhi, R., Wadher, B., Suryawanshi, N., & Selkar, S. (2017). Treatment of skin and soft tissue infections caused by Pseudomonas aeruginosa —A review of our experiences with citric acid over the past 20 years. Wound Medicine, 19, 5–9. 10.1016/j.wndm.2017.09.005

16. Nandini Shetty, Julian W Tang, J. A. (2009). Infectious Disease: Pathogenesis, Prevention and Case Studies. John Wiley & Sons.

17. Ng, C. Y., Huang, Y. H., Chu, C. F., Wu, T. C., & Liu, S. H. (2017). Risks for Staphylococcus aureus colonization in patients with psoriasis: a systematic review and meta-analysis. The British Journal of Dermatology, 177(4), 967–977. 10.1111/bjd.15366

18. Reiner, K. (2013). American Society for Microbiology, Catalase Test Protocol. American Society for Microbiology, November 2010, 1–9. http://www.microbelibrary.org/library/laboratory-test/3226-catalase-test-protocol

19. Sasson, G., Bai, A. D., Showler, A., Burry, L., Steinberg, M., Ricciuto, D. R., Fernandes, T., Chiu, A., Raybardhan, S., Science, M., Fernando, E., Morris, A. M., & Bell, C. M. (2017). Staphylococcus aureus bacteremia in immunosuppressed patients: a multicenter, retrospective cohort study. European Journal of Clinical Microbiology & Infectious Diseases : Official Publication of the European Society of Clinical Microbiology, 36(7), 1231–1241. 10.1007/s10096-017-2914-y

20. Sigudu, T. T., Oguttu, J. W., & Qekwana, D. N. (2024). Antimicrobial Resistance of Staphylococcus spp. from Human Specimens Submitted to Diagnostic Laboratories in South Africa, 2012-2017. Microorganisms, 12(9). 10.3390/microorganisms12091862

21. Termix. (2020, January). History and evolution of the hairbrush: burst of the XL hairbrush. Https://Termix.Net/Blog-Int/En/History-Evolution-Hairbrush-Burst-Xl-Brush/.

22. Type, I. (2015). hygiene): a grounded theory study . Personal grooming (beyond hygiene): a grounded theory study . A thesis submitted to meet the requirements of the University of Chester for the degree of Doctorate in Health and Social Care .

23. Ungphaiboon, S., Supavita, T., Singchangchai, P., Sungkarak, S., Rattanasuwan, P., & Itharat, A. (2005). Study on antioxidant and antimicrobial activities of turmeric clear liquid soap for wound treatment of HIV patients. Songklanakarin J. Sci. Technol., 27(April 2005), 569–578.

24. Vandini, A., Temmerman, R., Frabetti, A., Caselli, E., Antonioli, P., Balboni, P. G., Platano, D., Branchini, A., & Mazzacane, S. (2014). Hard Surface Biocontrol in Hospitals Using Microbial-Based Cleaning Products. PLoS ONE, 9(9), e108598. 10.1371/journal.pone.0108598

25. Wilkie, E. D., Alao, J. O., Sotala, T. T., & Oluduro, A. O. (2024). Molecular characterisation of virulence genes in bacterial pathogens from daycare centres in Ile-Ife, Nigeria: implications for infection control. BMC Infectious Diseases, 24(1), 1196. 10.1186/s12879-024-10095-8

26. Wilkinson, H. N., Stafford, A. R., Rudden, M., Rocha, N. D. C., Kidd, A. S., Iveson, S., Bell, A. L., Hart, J., Duarte, A., Frieling, J., Janssen, F., Röhrig, C., de Rooij, B., Ekhart, P. F., & Hardman, M. J. (2024). Selective Depletion of Staphylococcus aureus Restores the Skin Microbiome and Accelerates Tissue Repair after Injury. Journal of Investigative Dermatology, 144(8), 1865–1876.e3. 10.1016/j.jid.2024.01.018

